# Activity in the human superior colliculus associated with reaching for tactile targets

**DOI:** 10.1101/2022.09.06.506707

**Authors:** Nikhil G Prabhu, Marc Himmelbach

## Abstract

The superior colliculus (SC) plays a major role in orienting movements of eyes and the head and in the allocation of attention. Functions of the SC have been mostly investigated in animal models, including non-human primates. Differences in the SC’s anatomy and function between different species question extrapolations of these studies to humans without further validation. Few electrophysiological and neuroimaging studies in animal models and humans have reported an unexpected role of the SC in visually guided reaching movements. Using BOLD fMRI imaging, we sought to decipher if the SC is also active during reaching movements guided by tactile stimulation. Participants executed reaching movements to visual and tactile target positions. When contrasted against visual and tactile stimulation without reaching, we found increased SC activity with reaching not only for visual but also for tactile targets. We conclude that the SC’s involvement in reaching does not rely on visual inputs. It is also independent from a specific sensory modality. Our results indicate a general involvement of the human SC in upper limb reaching movements.

## Introduction

The Superior Colliculus (SC) is a mid-brain structure with a well-established role in orienting movements of the head, eyes, and arms (Gandhi and Katnani, 2011). Owing to decades of research on its contributions towards the generation of saccades and shifts of attention, the SC has been largely considered to be a primarily visually driven structure. The SC has been investigated for its role in visually guided arm movements in primates (Werner, 1993; Werner et al., 1997; Werner et al., 1997b) where it was shown that neurons in the SC were active either before, during, or after arm movements to visual stimuli. These observations were confirmed in humans by two fMRI studies on visually guided reaching (Linzenbold and Himmelbach, 2012; Himmelbach et al., 2013). Philipp and Hoffmann (2014) reported arm movements during electrical microstimulation of the SC in macaques, demonstrating that previously observed signal changes with upper limb reaching movements were not merely correlational signals but causally linked with movement execution. Cooper and McPeek (2021) recently reviewed evidence for the involvement of the SC in complex body and limb movements across species from lampreys to humans.

Beyond its dominant visual functions, the SC has long been known as a site processing multiple sensory modalities and performing multisensory integration (Meredith and Stein, 1983, 1985; Stein and Meredith, 1993). This raises the question whether previously observed signals during visually guided reaching were either visuomotor signals or general movement signals and independent from the sensory modality of target presentation. Targets can be presented through visual, auditory, or tactile information. Tactile target stimulations, which in contrast to auditory stimuli are not impeded by the background noise of an fMRI measurement, can be accurately localised by healthy humans on their own body (Weinstein, 1968; Schweizer et al., 2000; Azanõńand Soto-Faraco, 2008). Somatosensory body maps have been demonstrated in the SC of macaques and they are believed to be at least partially congruent with oculomotor maps (Ottes et al., 1986; Sparks and Nelson, 1987; Wallace et al., 1996; Nagy et al., 2006).

We measured BOLD fMRI during reaching movements guided by tactile target information in five participants. Reaching to tactile targets was compared with visually guided reaching. The participants placed the palm of their left hand on a palm-rest above their waist (Figure 5 A, B). Tactile stimulation was applied at the fingertips with pneumatic stimulators (Figure 5 C). Visual targets were presented through fibre optic cables terminating above each fingertip (Figure 5 A, B). The participants reached with their right index finger from a start button at their chest to the left hand’s fingertip at the target location. Reaching movements in either sensory modality were compared with corresponding sensory control conditions. In sensory control conditions, we presented the same visual and tactile target stimuli. The participants pressed the start button if a rare oddball stimulus occurred. We expected signal increases during reaching with both target modalities at the SC signal peak location that was observed in earlier studies on visually guided reaching (Linzenbold and Himmelbach, 2012; Himmelbach et al., 2013).

## Results

### Behavioural data

Reaching to visual targets, the participants showed a mean reaction time of 0.38 s with a standard deviation (SD) of 0.03 s. The mean reaction time for reaching to tactile targets was 0.43 s (SD 0.06 s). The mean duration of visually guided reaching was 1.53 s (SD 0.18), with 1.65 s (SD 0.16 s) for reaching to tactile targets. The participants detected 54 to 56 oddball targets in both the visual control condition and tactile control condition out of a total of 56 oddball targets. The mean reaction time for button presses to oddball stimuli in the visual control condition was 0.54 s (SD 0.18 s) and 0.68 s (SD 0.17 s) for tactile oddballs. The average number of saccades per run varied between 0 and 7 out of a total of 112 trials in subjects 1,2,3 and 5. For the 4th subject it varied between 0 and 48. These numbers included all runs available for analysis i.e. those that were made of reaching conditions only, sensory conditions only and mixed runs. In reaching conditions the number of saccades per run varied between 0 and 2 and in the sensory control conditions between 0 and 7 out of a total of 112 trials in the four subjects mentioned above. In the 4th subject however the number of saccades per run in reaching conditions varied between 0 to 48, and in the sensory control conditions between 3 and 43 out of a total of 112 trials.

### BOLD fMRI results

We focused our analysis on the VOI (voxel-of-interest) −6, −28, −6 in MNI (Montreal Neurological Institute) co-ordinates. We chose this voxel location because it was the peak signal location in GLM (General Linear Model) group analyses of SC signals during visually guided reaching (Himmelbach and Linzenbold, 2012), confirmed in a following study on visually guided anti-reaching (Himmelbach et al. 2013). The VOI is located in the deep left SC, where we expected a motor signal corresponding to the contralateral, right reaching arm. Additionally, we analysed signals at the corresponding ipsilateral location in the right SC (6, −28, −6). We estimated a GLM using SPM12 (Statistical Parametric Mapping, version 12) with four experimental conditions, button presses, and visual cues indicating conditions as predictors and extracted contrast estimates with 90% confidence intervals as measures of activity at the VOI (Figure 1, Figure 3). Where the confidence intervals did not overlap with zero, we would consider this as a significant finding. The contrast estimates during both reaching modalities were significantly positive in 5 out of 5 participants (Figure 1). In the sensory control conditions the estimates’ confidence intervals mostly overlapped with zero or the estimates were negative relative to the baseline (Figure 1). The estimates from the differential contrasts between reaching and sensory control conditions confirmed this observation. For visual targets, the difference was significantly positive in 4 out of 5 participants (Fig. 2; VM>VS). The same number of participants showed a significantly positive difference for tactile stimuli (Fig. 2; TM>TS). Where the differential signal was not significant, it was still positive (Figure 1; Subject 4 VM>VS and Subject 2 TM>TS). The VM>TM comparison shows the signal levels to be around zero in 4 out of 5 subjects and negative in one subject (subject 1) (Figure 1). This indicates that the signal levels for reaching to visual and reaching to tactile targets were equivalent, except for the 1st subject where the tactile reaching condition shows a slightly increased signal level compared to visually guided reaching.

**Figure 1:**
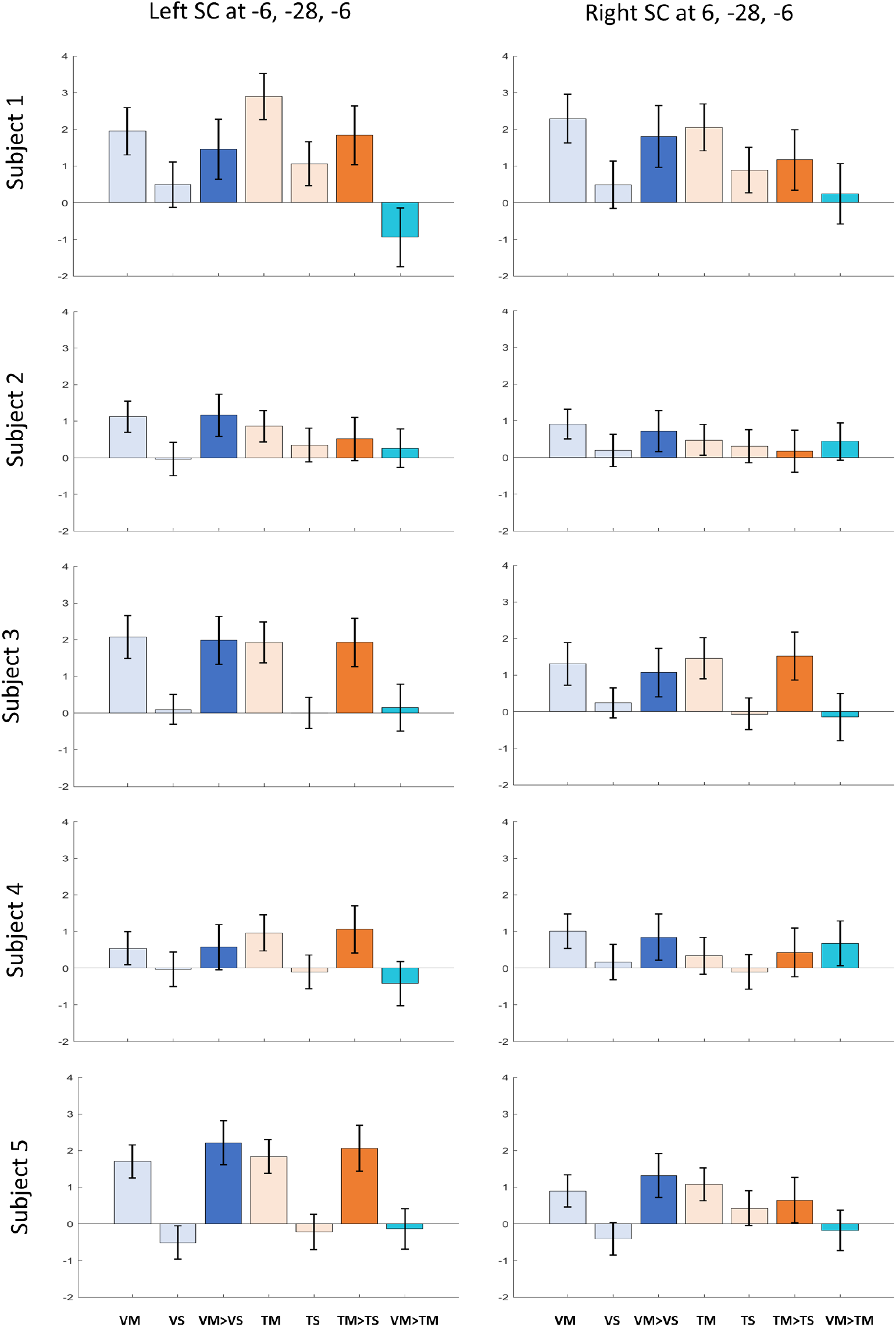
Contrast estimates from the conditions of the experiment at −6, −28, −6 indicate higher signal contributions from the motor conditions when compared to their respective controls. Contrast estimates for the two motor conditions VM and TM and the two control conditions VS, TS along with the differential estimates VMVS, TMTS and VMTM. The black bars indicate 90% confidence intervals (CI).

**Figure 2:**
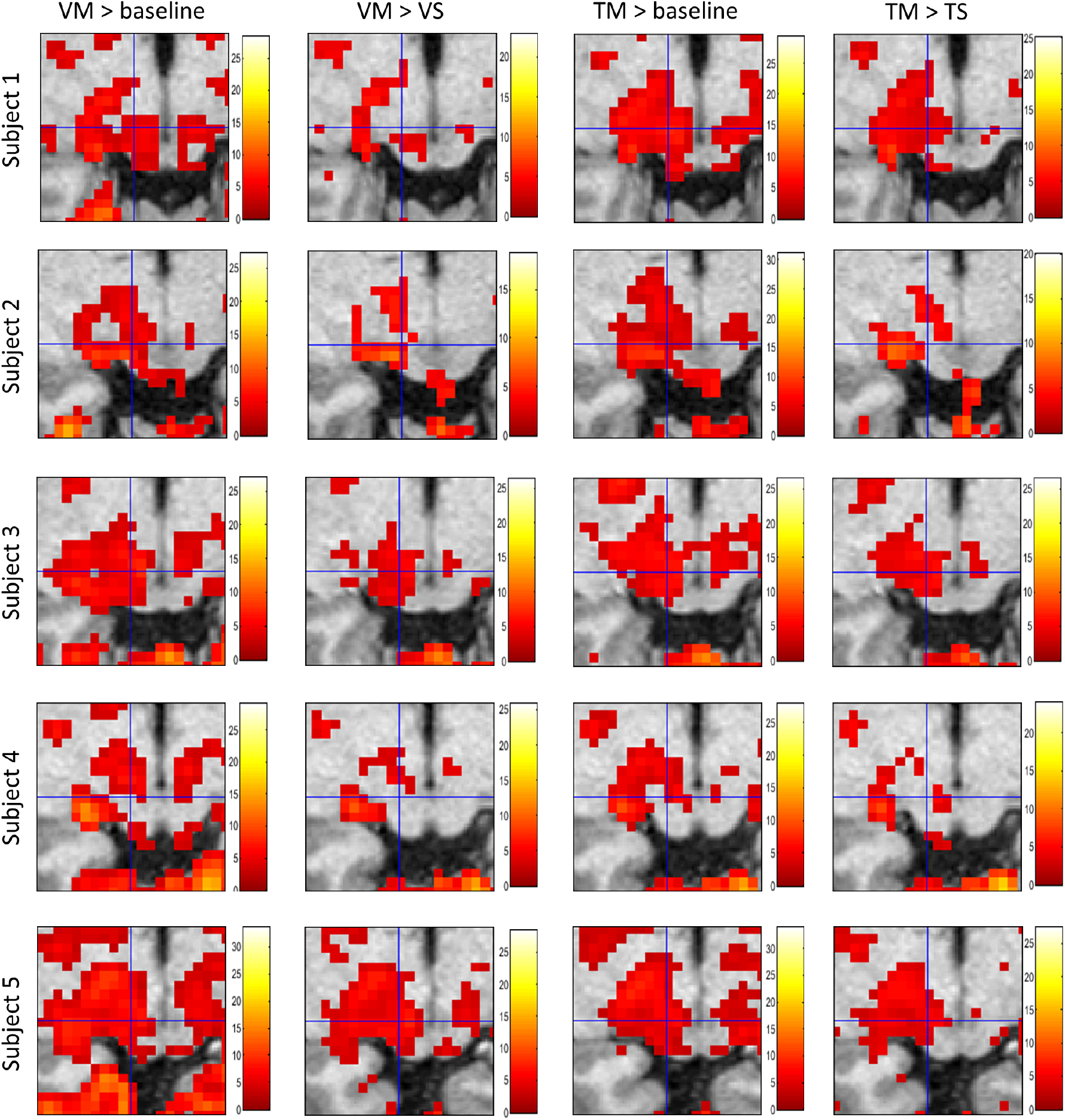
VM and TM show a robust signal when compared to baseline and control conditions. The figures show a transverse section of the SC (neurological convention) centred at −6, −28, −6, indicated by the crosshair. Columns show significant signal increases (*p* < 0.001, uncorrected) for the contrasts - VM>baseline (A), VM>VS (B), TM>baseline (C) and TM>TS (D), with each row corresponding to a subject. The colorbar next to each figure denotes the range of T values.

**Figure 3:**
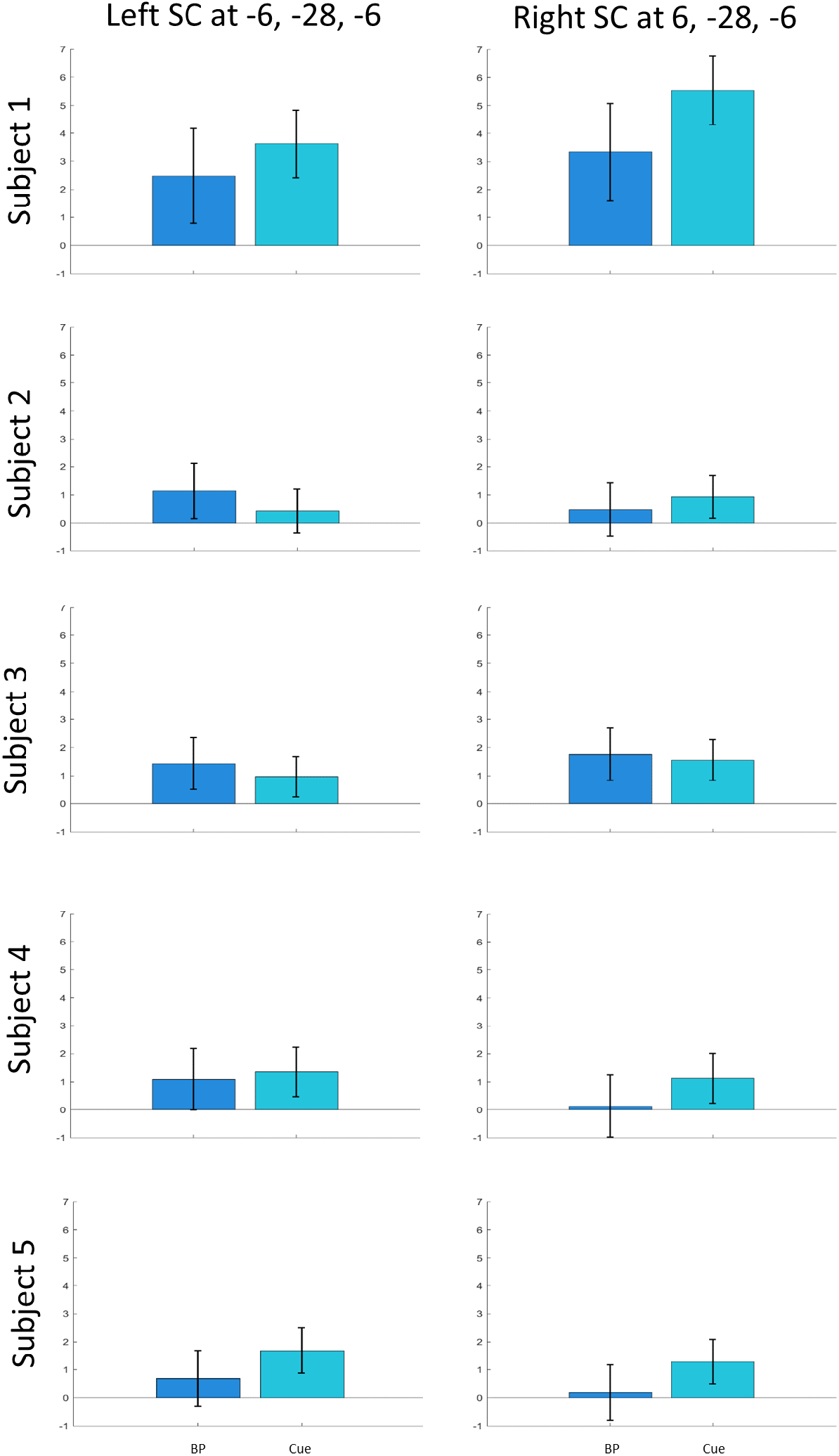
Contrast estimates from the button press and cue conditions show a positive signal contribution at the VOI −6, −28, −6. Signal contributions plotted at the VOI and its corresponding location in the right SC from the button press (BP) and cue conditions. The black bars indicate 90% confidence intervals (CI).

To contextualize the VOI results above, we plotted voxel-wise GLM results on transverse sections of the SC from all participants with crosshairs centered on the VOI in Figure 2. We thresholded voxels with *p* < 0.001 uncorrected and observed suprathreshold clusters at or next to the VOI in both reaching conditions versus baseline in all 5 subjects (Figure 2, 1st and 3rd column). Also for the contrasts between reaching and sensory control conditions, we detected voxels and clusters at or immediately next to the VOI location (Figure 2, 2nd and 4th column). We found no significant activations for the contrasts of the sensory control conditions against the baseline. Finally, we inspected signal contributions from rare button presses (1 in 8 subsequent trials) in the sensory control conditions and from visual cues preceding the experimental condition blocks. The signal from button presses was mostly positive at the VOI (Figure 3). The signal contribution from cues was mostly positive at the VOI and at the corresponding location in the right SC.

Signal time courses of both motor conditions showed a consistent increase temporally aligned with the duration of a block of trials at the VOI in all 5 participants (Figure 4 A). The signal is seen plateauing in the middle of the block and begins to dip towards the end of the block. The time courses of the sensory control conditions do not show systematic changes throughout the block across the participants. The time courses for visual and tactile conditions followed a similar pattern without systematic differences. Based on the GLM voxel-wise results of the comparisons between motor and sensory control conditions reported above (Figure 2), we identified the most and the least significant voxel, immediately next to the VOI. While time courses at the most significant voxels showed a pattern very similar to time courses from the VOI (Figure 4 B), time courses from the least significant voxels do not show any pattern, apart from a slight positive increase in the visual reaching condition at the start of the block in 4 of the 5 subjects (Figure 4 C).

**Figure 4:**
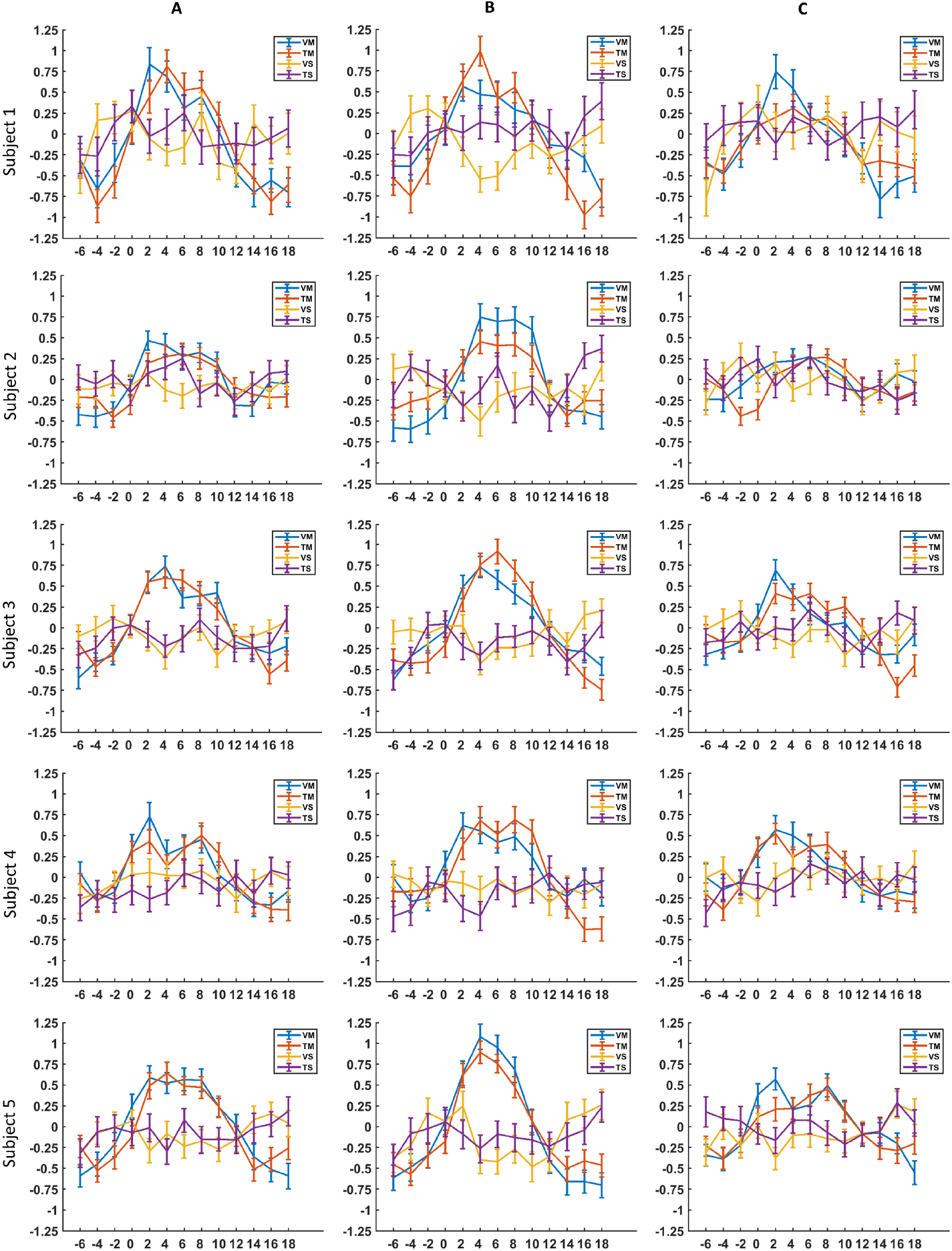
Figure showing averaged time courses from the four conditions at the VOI, most significant, and least significant voxel, from the 27 voxel grid in the SC. A – VOI (voxel-of-interest), −6, −28, −6; B, C – voxels with the least p-value (most significant result) and highest p-value (least significant result) from the 27 voxel grid in the SC for each condition. The four conditions (VM, VS, TM and TS) are labelled in the legend in the upper right corner. The error bars show standard error (SE).

## Discussion

We detected signal increases in the SC for reaching movements to tactile targets at the expected location in the deep SC in all five participants of our study. These signal increases were equivalent to signal changes during reaching to visual targets and exceeded signals due to sensory stimulation alone in four out of five participants. These results confirmed that the SC represents a subcortical node in a general sensorimotor network as shown by previous studies in humans (Linzenbold and Himmelbach, 2012; Himmelbach et al., 2013) and non-human primates (Werner, 1993; Werner et al., 1997b; Lünenburger et al., 2001). Our current findings demonstrated that this involvement is not constrained to the integration of visual input.

As time progresses, there is greater evidence that the SC is involved in functions that traverse beyond visual detection, orientation movements of the eyes and head, and saccade generation. The functions of the SC/tectum is more or less conserved across vertebrates with its involvement in several complex behaviours. There are several examples that reflect this - the murine SC is known to take part in contradictory movement patterns such as approach and aversion along with other complex behaviours such as defensive posturing, aggression etc. (Olds and Olds 1963; Valenstein 1965; Panksepp 1971; Waldbillig 1975; Imperato and Di Chiara 1981; McHaffie and Stein 1982; Weldon et al., 1983). The zebrafish tectum based on the inputs it receives likely carries information on modalities like auditory, vestibular and potential water flow (Isa et. al 2021). Rattlesnakes combine thermal and visual information to identify prey in complex environments and in darkness (Hartline et al., 1978; Newman and Hartline, 1981; Standford and Hartline, 1984). The auditory spatial maps in barn owls allows it to hunt when it is dark (Knudsen and Konishi, 1978; Knudsen 1982; Konishi 1993; Carr and Konishi, 1990). Along with this, the mammalian SC such as in mice, cats, monkeys and humans is known to play a role in behaviours such as forelimb reaching movements, target selection and decision-making (Basso and May, 2017; Cooper and McPeek, 2021; Isa et al., 2021). Taken together, these studies indicate that the SC is a multisensory integration center that is involved in the generation of complex behaviours. The current study was conceived with the intention to expand this knowledge further and probe a specific sensory modality - tactile stimulation, for its role in guiding reaching movements in the SC.

Although, previous studies have looked at processing of tactile signals in the SC with respect to saccades (Stein and Meredith, 1993; Groh and Sparks, 1996a, 1996b, 1996c), they have not gone as far as to study if it in some way relates to reaching movements. The only study that related tactile information to reaching is the one by Nagy et al. (2006) which suggests that the SC may be involved in interaction with objects. In this study, neuronal activity was seen in the SC when the hand of a monkey came in contact with an object, with the activity modulated by increased force. The question of whether processing of somatosensory signals in the SC helps in the execution of reaching movements had not yet been probed. All the other studies pertaining to the SC that used somatosensory stimulation have examined it in the context of execution of saccades or orienting movements of the head. As far as we know the only studies indicating that a sensory modality could help in homing in on targets without the help of vision is hunting in the dark based on auditory spatial maps in the SC of barn owls (Knudsen and Konishi, 1978; Knudsen 1982; Konishi 1993; Carr and Konishi, 1990). Nagy et al. (2006) speculate that similar to the monitoring of errors during fixation in fixation neurons of the SC (Krauzlis et. al, 2017), the somato-sensory neurons that they found in the SC might be involved in monitoring small changes in muscle forces that are needed to interact effectively with a target that is reached. In addition, we propose that reaching to tactile targets could help in complete darkness or when a faster response is necessary than the combination of tactile stimulation and subsequent foveation, before a reaching movement is made. For example, when an insect is felt on the skin, the first reaction is to brush it away as soon as possible. This is generally followed by foveation for recognition of the insect.

The SC is known to possess auditory and somatosensory maps which are aligned with visual and motor maps (Drager and Hubel, 1975; Chalupa and Rhoades, 1977; Stein and Meredith, 1993; Ghose et al., 2014). Somatosensory maps have been proposed to have been organized relative to how body parts would be seen from the eye, hinting that information from these maps are probably being converted to retinotopic co-ordinates (Drager and Hubel, 1975). Groh and Sparks (1996) found that almost all the cells in the SC that responded to saccades for visual targets also did so for saccades to tactile targets. These findings are in agreement with our results which show no statistical difference in signals between visually guided reaching and reaching guided by tactile stimulation. Although with our study, it is not possible to prove if the tactile targets were represented in retinotopic co-ordinates, we were able to prove that visual information is not necessary to execute reaching movements in humans.

Some of the first studies on reaching in primates were those by Werner (1993) and Werner et al. (1997). In the latter study, it was shown that most of the cells probed were tuned to the direction of reaching, called the reach movement field and that there was a correlation between the onset of activity in these cells and the EMG (electro-myogram) of the muscles involved in reaching movements. In a later study Stuphorn et al. (2000) examined this spatial tuning in more detail and found that there were two sets of reach neurons in the SC - one population of cells present in the intermediate and deeper layers with a presence of activity for arm movements corresponding only to the retinal location of the targets called gaze-dependent neurons and a second population of cells in the deeper layers of the SC and reticular formation that encoded the direction of arm movements irrespective of the location of the reach target on the retina. All these studies found that the preferred direction of cells did not depend on the organization of visual and saccade-related cells in the SC. Importantly, all of these studies also found that reach related-activity depended on the position of the target but not on the trajectory of the reach movement nor on the EMG of the muscles that were involved in the reach. This in turn indicates that the reach-related neurons provide abstract signals relating to the location of the reach goals than to the kinematics of reaching itself (Cooper and McPeek, 2021). Nevertheless, Cooper and McPeek (2021) also argue that reach-related pathways in the SC could be involved in correction of reaching movements, eye-hand co-ordination and error correction after targets have been contacted and warrant more studies for further elucidation.

Although our results indicate that saccades are not correlated with reaching movements, an argument stating that microsaccades executed during reaching movements could have caused the activation in the SC, is plausible. An earlier study in macaques by Reyes-Puerta et al. (2010) indicated that ‘reach’ neurons in the SC show increased activity in reaching trials where trials with microsaccades were excluded. This means that although there is a possibility that a part of the signal could be owed to microsaccades correlated with reaching movements, it is unlikely that the signal is completely due to them.

The experiment teases apart activity in the SC towards reaching movements corresponding to visual and tactile targets from that which could have been expected from sensory stimulation alone. Although it is established that a representation for visual or tactile stimulation on their own exists in the SC, we did not see it in our experiment. This could be due to the large distance (9°) between the centre of the four visual targets and the fixation LED. Chen et al. (2019) have shown that responses to visual stimuli in the SC are tightly linked with foveation. Hence, there could have been an under-representation of these peripheral targets. The tactile stimulation was probably too weak and lasted for a very short time for the corresponding activity to be sensitive enough for detection with fMRI in the SC.

An earlier study from the Himmelbach group (Linzenbold and Himmelbach, 2012) showed a strong lateralization in signals from left and right hand reaching. In comparison, responses to reaching in the SC from our study are not clearly lateralised. This might have to do with an entirely different configuration of reaching trials in both the experiments with the main difference being that reaching movements were directed towards the contralateral arm in our experiment. This is similar to the gaze-independent neurons in the SC responding more strongly to contralateral arm position (Lünenburger et al., 2001). Besides, Werner et al. (1997) mention the presence of ipsilateral and bilateral neurons in the SC in addition to contralateral ones. Stuphorn et al. (2000) see that a subset of deep SC neurons called the gaze-dependent neurons show activity in response to reaching by both contralateral and ipsilateral arms. Particularly, Nagy et al. (2006) showed contact-related responses for both ipsilateral and contralateral arm movements. In contrast to the group analysis in Linzenbold and Himmelbach (2012), we conducted single-case analyses in the current study. In our single-case analyses, we also find a relative lateralization in the activity estimates in most participants (see Figure 2). The difference between stronger contrasts between contra- and ipsilateral reaching in Linzenbold and Himmelbach (2012) and the more bilateral signals in the indvidual datasets reported here might, partially, be due to different measurement and analysis strategies.

In conclusion, the SC seems to integrate sensory input from a range of modalities towards a functional contribution to motor responses in humans. This link between the SC and motor responses of the upper limb is in good agreement with observations of motor control signals in the SC or tectum across multiple, non-human species.

## Materials and methods

### Subject details

We conducted the study using 3T BOLD fMRI on 5 subjects (5 females, age range 22-28, all right-handed) with normal or corrected-to-normal visual acuity. Each subject performed the experiment four times, one session per day. We excluded the first two sessions from the first subject for non-conformity with the rest of the measurements, since they were measured with the online motion correction of the scanner software. We conducted the experiments with the approval of the local ethical committee and as per the ethical standards established by the 1964 declaration of Helsinki. We obtained informed consent from all the subjects.

### Experimental setup

Subjects performed the experiments in complete darkness. We ensured no light penetrated the scanner room by covering windows, scanner displays, and the opening of the scanner bore with black opaque sheets. Subjects rested the palm of their left hand on a palm-rest (Figure 5 B) with four slits shaped according to the four fingers used for the experiment (excluding the thumb). The palm-rest in turn rested on a stand placed across the waist of the subjects (Figure 5 A). Pneumatic stimulators (Figure 5 C) were placed in a notch on the uppermost part of the slits in the palm-rest making contact with the respective fingertips. Changes in air pressure brought about by a compressor regulated the diaphragms of the pneumatic stimulators, causing tactile stimulation on fingertips. Fibre optic cables carried light from green and red LEDs into holes drilled above each of the four finger slits on the palm rest (Figure 5 A, B). 8 fibre optic cable endings reached the fingertips: 2 per fingertip, 1 of each colour, hence, making up four targets. Each target would glow red or green depending on the trial type. The white LED formed the fixation point and was attached to the scanner bore. The targets were placed at a distance of 9°away from the fixation point, and at a varying distance of 4°, 3°and 2°between them (from the index finger to the little finger, respectively) owing to the angle subtended between the four fingers. Subjects looked at the fixation point directly, eliminating the need for a mirror. We placed cushions specially manufactured for fMRI research (NoMoCo Pillow, Inc., La Jolla, USA) into the head-coil of the scanner, immobilising the head of the subjects. We programmed a microcontroller - ‘Mbed LPC1768’ (Arm Limited.) to control target LEDs, and air pressure regulating valves connected to pneumatic stimulators. In addition, the microcontroller was also programmed to collect button press timings, display experimental events on screen, and write all event timings to a file. We used two MR compatible cameras (MRC systems GmbH.) that operated at 30Hz: one to check fixation and the other to monitor reaching movements to targets. The cameras were in turn connected to a frame grabber (Matrox Imaging) which was controlled via MATLAB (version R2018a, The MathWorks Inc., Natick, MA, USA) for video recording. The first TTL pulse from the scanner triggered the microcontroller to execute the experimental paradigm and the frame grabber to start video capture in MATLAB.

**Figure 5:**
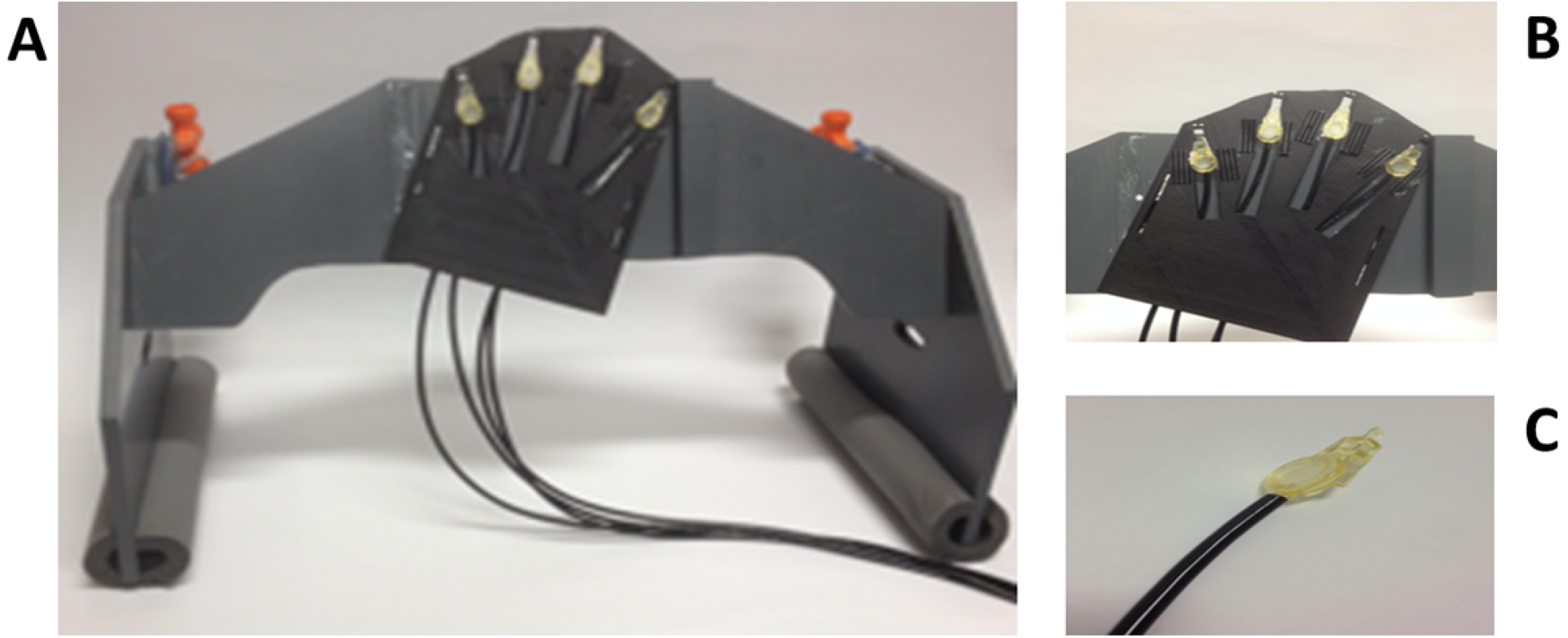
Apparatus. A - The stand onto which the palm rest was attached. B – palm rest with attached pneumatic stimulators and air tubes. C – a single pneumatic stimulator.

### Experimental paradigm

Each experimental session consisted of four conditions – VM (visuo-motor); TM (tactile-motor); VS (visual sensory); and TS (tactile sensory); put to-gether in a factorial design, resulting in a combination of two conditions in each of the four runs. The four runs thus formed comprised of the following conditions: run 1 - VM, TM; run 2 - VM, VS; run 3 – VS, TS and run 4 - TM, TS. The conditions that required a reaching response: VM and TM were our conditions of interest while the conditions that required sensory discrimination: VS and TS, formed the controls. Green and red LEDs formed the targets in the visual conditions, and single or double tactile stimulation formed the targets in the ‘tactile’ conditions. Subjects were required to fixate on the central white LED above the targets throughout the experiment. The central fixation light turned on with the receipt of the first TTL pulse from the scanner marking the start of a run. All runs started with an initial fixation period of 12s. The two conditions forming a run alternated as blocks of four targets each, with each block adding up to 8.8 seconds (stimulus time of 0.2s and inter-stimulus interval of 2s). A 12s baseline period followed each block apart from the set of blocks where the sensory modality was the same i.e. the task to be performed in the impending block was not obvious. Such blocks were preceded by a cue (blink of the fixation light) within the 12s baseline period, 3 seconds before the start of the block. For example, tactile stimulation (TS) and tactile reaching (TM) blocks were preceded by one blink and two blinks of the cue respectively. Both the blocks repeated 14 times each, adding up to around 10 minutes of scanning time for a run. Each of the four targets in a block appeared in a pseudo-randomised manner with most of the tactile stimulation being single stimulation, and visual targets being green in colour depending on the respective block. One in eight targets was an oddball: either red in colour or a double tactile stimulation depending on the condition block they belonged to. In the sensory conditions, subjects were required to respond to the oddball (red target or double tactile stimulation) with a button press. This resulted in 28 button presses per session, totalling 112 button press responses per subject with all four sessions combined. In the reaching conditions, subjects had to make a reaching movement towards a target with their right hand irrespective of whether it was green or red in colour, or whether it was a single or double tactile stimulation i.e. irrespective of whether it was a regular target or an oddball. Only the forearm was used to make reaching movements while the rest of the arm fixed to the scanner bed. This was done to reduce movements of the head while executing reaching movements. The total number of block repetitions per subject with all the four sessions combined were 448 (112 block repetitions from each of the four conditions VM, TM, VS and TS) with the exception of the first subject with 224 repetitions (since two sessions were excluded). To register the time taken to make the movement, subjects continuously pressed a button placed on their chest, releasing it only to make reaching movements in response to peripheral targets and returning to it soon after. In the sensory discrimination conditions, responses were recorded when subjects pressed a second button with the index finger of their right hand.

### fMRI data acquisition

We acquired the images on a Siemens Prisma 3 Tesla MRI scanner with a 32-channel head coil. 300 T2* weighted functional images (echo-planar imaging (EPI) sequence) were acquired per run with a slice thickness of 2mm, TR=2s, TE=35ms, flip angle of 80°and FOV - 192 × 192 mm (96 × 96 matrix). Each functional image consisted of 23 slices in a transversal orientation covering structures of our interest: the SC and the primary visual cortex (V1). We also acquired high resolution T1 weighted structural images once per subject (MP-RAGE sequence, slice thickness of 0.8 mm, TR=2.4s, TE=2.22ms, flip angle of 8°and FOV – 240 × 256 mm (300 × 320 matrix). A whole-brain EPI image was acquired in each of the four sessions of a subject with the same parameters as above except for the number of slices (72) and TR (6.1s). The whole-brain EPI images were acquired to ensure good co-registration between the functional and whole-brain images (details in the pre-processing section below).

### Analysis

#### Behavioral data

Compliance with task instructions was ensured through online monitoring of the hand and eye video recordings. We calculated reaction times and movement duration from the press and release of a button at the right hand’s start location. Due to technical issues we lost some eye video recordings for a detailed, offline analysis of gaze fixation. 3 of 8 runs were still available in subject 1, 7/16 runs in subject 2, 16/16 of subject 3, 16/16 of subject 16, and 4/16 in subject 5 (46 runs out of a total of 72). We extracted eye traces from video recordings with DeepLabCut (Mathis et al., 2018). A threshold of 2°(visual angle) was applied for extracting saccades from these eye traces using Python (version 3.7.10, Python Software Foundation, https://www.python.org). Synchronisation errors in the frame grabber made the timing of some videos unreliable. Some random frames were randomly lost in the available videos. Therefore, a reliable temporal analysis of these videos was not possible hence, we analysed the total number of saccades per run. We used runs with only reaching and sensory conditions - 2 runs out of 4 in each session to calculate the number of saccades in reaching and sensory runs which we have reported separately.

#### Pre-processing of fMRI data

We used SPM12 (Statistical Parametric Mapping, version 12; Wellcome Trust Centre for Neuroimaging, London, UK) implemented in MATLAB (version R2018a, The MathWorks Inc., Natick, MA, USA) to pre-process the images and later perform a GLM analysis. The first 5 images at the start of all runs were deleted to allow signals to reach a steady state. This was followed by a re-alignment procedure (as defined in SPM) on the rest of the images. We then co-registered the mean functional image (resulting from re-alignment) with the whole-brain EPI image followed by a coregistration of the whole-brain EPI image with the anatomical image (T1). This was performed for each of the four sessions in a subject, separately, to achieve good registration between the functional images (partial volumes) and the anatomical images. We confirmed the co-registration by matching the functional images with the anatomical scans in each session. We then normalised the anatomical and functional images to the MNI152 (Montreal Neurological Institute) template. We inspected the co-registration in each subject and manually adjusted the outcome of the automatic coregistration procedure resulting in alignment of all the functional images from the four sessions to one anatomical image (from the first session), in each subject. This was followed by smoothing of all functional images with a FWHM (Full Width Half Maximum) Gaussian kernel of 3mm.

#### BOLD fMRI data analysis

We used SPM12 for a first-level analysis pooling all the data from the four sessions in each subject. We modified the HRF (Haemodynamic Response Function) parameters to use an onset-to-peak time of 4s which is considered apt for the SC (Wall et al., 2009). A mask of the CSF space adjoining the SC was created for all subjects using MRICron (NITRC, University of South Carolina, Columbia, SC, USA). A mean time course across all functional images per subject was obtained from the mask using Marsbar (http://marsbar.sourceforge.net). This mean time course, along-with motion parameters derived from the re-alignment procedure in the pre-processing stage, were used as regressors of no-interest. We used the button press condition and cues as additional regressors of interest. We used movement onset times as onsets in the GLM analysis for the reaching conditions. In the sensory discrimination conditions, button press onsets were used for oddball stimuli, while onsets for non-oddball stimuli and cues were derived by an addition of mean reaction times from the reaching conditions to the onset times of stimulus presentation. This ensured coherence in onsets between conditions. The duration of each of the events was set to 0. The results were analysed centred at our VOI (voxel-of-interest) −6, −28, −6 in MNI (Montreal Neurological Institute) co-ordinates.

To examine the raw data acquired from the experiment in our primary VOI (−6, −28, −6) and the voxels surrounding it in the deep layers of the SC, we created a 3×3×3 grid (forming a cube of 27 voxels) centred around the primary voxel. Next, we extracted the raw time course for each voxel with a high-pass filter of 128 hz. We de-meaned the time courses and used the same regressors of no-interest as in the GLM analysis (motion parameters + mean time course of the CSF mask). We interpolated the time course to 0.1 seconds from the 2s TR (Repetition time) and then resampled the data back to 2s starting from stimulus or movement onsets, as applicable, in each of the four conditions. Time courses from each of the four conditions were then averaged across all repetitions to give rise to one time course per condition, per voxel, in each subject. These time courses were then used to examine changes during the block across subjects for each condition.

## Acknowledgments

We thank Dr. Michael Erb, Department for Biomedical Magnetic Resonance, University Hospital of Tübingen for his assistance with fMRI data acquisition protocols and related troubleshooting. We thank Prof. Christoph Braun and Mr. Juergen Dax, MEG center, University Hospital, Tuebin- gen for providing tactile stimulators and for assistance with building the tactile stimulation rig. This project was supported by a grant awarded by the DFG to MH (DFG Hi 1371-1-2)

## References

Alex Meredith, M., and Stein, B.E. (1983). Interactions among converging sensory inputs in the superior colliculus. Science (80-.). 221, 389–391. https://doi.org/10.1126/science.6867718.

Azañón, E., and Soto-Faraco, S. (2008). Spatial remapping of tactile events. Commun. Integr. Biol. 1, 45–46. https://doi.org/10.4161/cib.1.1.6724.

Basso, M.A., and May, P.J. (2017). Circuits for Action and Cognition: A View from the Superior Colliculus. Annu. Rev. Vis. Sci. 3, annurev-vision-102016-061234. https://doi.org/10.1146/annurev-vision-102016-061234.

Carr, C.E., and Konishi, M. (1990). A circuit for detection of interau-ral time differences in the brain stem of the barn owl. J. Neurosci. 10, 3227–3246. https://doi.org/10.1523/jneurosci.10-10-03227.1990.

Chalupa, L.M., and Rhoades, R.W. (1977). Responses of visual, somatosen-sory, and auditory neurones in the golden hamster’s superior colliculus. J. Physiol. 270, 595–626. https://doi.org/10.1113/jphysiol.1977.sp011971.

Chen, C.Y., Hoffmann, K.P., Distler, C., and Hafed, Z.M. (2019). The Foveal Visual Representation of the Primate Superior Colliculus. Curr. Biol. 29, 2109-2119.e7. https://doi.org/10.1016/j.cub.2019.05.040.

Cooper, B., and McPeek, R.M. (2021). Role of the Superior Colliculus in Guiding Movements Not Made by the Eyes. Annu. Rev. Vis. Sci. 7, 279–300. https://doi.org/10.1146/annurev-vision-012521-102314.

Drager, U.C., and Hubel, D.H. (1975). Responses to visual stimulation and relationship between visual, auditory, and somatosensory inputs in mouse superior colliculus. J. Neurophysiol. 38, 690–713. https://doi.org/10.1152/jn.1975.38.3.690.

Foley, H., and Matlin, M. (2021). The Skin Senses. Sensat. Percept. 344–370. https://doi.org/10.4324/9781315665061-18.

Gandhi, N.J., and Katnani, H.A. (2011). Motor functions of the superior colliculus. Annu. Rev. Neurosci. 34, 205–231. https://doi.org/10.1146/annurev-neuro-061010-113728.

Ghose, D., Maier, A., Nidiffer, A., and Wallace, M.T. (2014). Multisensory response modulation in the superficial layers of the superior colliculus. J. Neurosci. 34, 4332–4344. https://doi.org/10.1523/JNEUROSCI.3004-13.2014.

Groh, J.M., and Sparks, D.L. (1996a). Saccades to somatosensory targets. I. behavioral characteristics. J. Neurophysiol. 75, 412–427. https://doi.org/10.1152/jn.1996.75.1.412.

Groh, J.M., and Sparks, D.L. (1996b). Saccades to somatosensory targets. II. Motor convergence in primate superior colliculus. J. Neurophysiol. 75, 428–438. https://doi.org/10.1152/jn.1996.75.1.428.

Groh, J.M., and Sparks, D.L. (1996c). Saccades to somatosensory targets. III. eye-position-dependent somatosensory activity in primate superior colliculus. J. Neurophysiol. 75, 439–453.

Hartline, P.H., Kass, L., and Loop, M.S. (1978). Merging of modalities in the optic tectum: Infrared and visual integration in rattlesnakes. Science (80-.). 199, 1225–1229. https://doi.org/10.1126/science.628839.

Himmelbach, M., Linzenbold, W., and Ilg, U.J. (2013). Dissociation of reach-related and visual signals in the human superior colliculus. Neuroimage 82, 61–67. https://doi.org/10.1016/j.neuroimage.2013.05.101.

Imperato, A., and Di Chiara, G. (1981). Behavioural effects of GABA-agonists and antagonists infused in the mesencephalic reticular formation - deep layers of superior colliculus. Brain Res. 224, 185–194. https://doi.org/10.1016/0006-8993(81)91131-8.

Isa, T., Marquez-Legorreta, E., Grillner, S., and Scott, E.K. (2021). The tectum/superior colliculus as the vertebrate solution for spatial sensory integration and action. Curr. Biol. 31, R741–R762. https://doi.org/10.1016/j.cub.2021.04.001.

Knudsen, E.I. (1982). Auditory and visual maps of space in the optic tectum of the owl. J. Neurosci. 2, 1177–1194. https://doi.org/10.1523/jneurosci.02-09-01177.1982.

Knudsen, E.I., and Konishi, M. (1978). A neural map of auditory space in the owl. Science (80-.). 200, 795–797. https://doi.org/10.1126/science.644324.

Krauzlis, R.J., Goffart, L., and Hafed, Z.M. (2017). Neuronal control of fixation and fixational eye movements. Philos. Trans. R. Soc. B Biol. Sci. 372, 20160205. https://doi.org/10.1098/rstb.2016.0205.

Linzenbold, W., and Himmelbach, M. (2012). Signals from the deep: Reach-related activity in the human superior colliculus. J. Neurosci. 32, 13881–13888. https://doi.org/10.1523/JNEUROSCI.0619-12.2012.

Lünenburger, L., Kleiser, R., Stuphorn, V., Miller, L.E., and Hoffmann, K.P. (2001). A possible role of the superior colliculus in eye-hand coordination. Prog. Brain Res. 134, 109–125. https://doi.org/10.1016/S0079-6123(01)34009-8.

Mathis, A., Mamidanna, P., Cury, K.M., Abe, T., Murthy, V.N., Mathis, M.W., and Bethge, M. (2018). DeepLabCut: markerless pose estimation of user-defined body parts with deep learning. Nat. Neurosci. 21, 1281–1289. https://doi.org/10.1038/s41593-018-0209-y.

McHaffie, J.G., and Stein, B.E. (1982). Eye movements evoked by electrical stimulation in the superior colliculus of rats and hamsters. Brain Res. 247, 243–253. https://doi.org/10.1016/0006-8993(82)91249-5.

Meredith, M.A., and Stein, B.E. (1985). Descending efferents from the superior colliculus relay integrated multisensory information. Science (80-.). 227, 657–659. https://doi.org/10.1126/science.3969558.

Nagy, A., Kruse, W., Rottmann, S., Dannenberg, S., and Hoffmann, K.P. (2006). Somatosensory-Motor Neuronal Activity in the Superior Colliculus of the Primate. Neuron 52, 525–534. https://doi.org/10.1016/j.neuron.2006.08.010.

Newman, E.A., and Hartline, P.H. (1981). Integration of visual and infrared information in bimodal neurons of the rattlesnake optic tectum. Science (80-.). 213, 789–791. https://doi.org/10.1126/science.7256281.

Olds, M.E., and Olds, J. (1963). Approach-avoidance analysis of rat diencephalon. J. Comp. Neurol. 120, 259–295. https://doi.org/10.1002/cne.901200206.

Ottes, F.P., Gisbergen, J.A.M. van, and Eggermont, J.J. (1986). Visuomotor Colliculus : Fields of the Superior a Quantitative Model. 26, 857–873.

Panksepp, J. (1971). Aggression elicited by electrical stimulation of the hypothalamus in albino rats. Physiol. Behav. 6, 321–329. https://doi.org/10.1016/0031-9384(71)90163-6.

Philipp, R., and Hoffmann, K.P. (2014). Arm movements induced by electrical microstimulation in the superior colliculus of the macaque monkey. J. Neurosci. 34, 3350–3363. https://doi.org/10.1523/JNEUROSCI.0443-13.2014.

Reyes-Puerta, V., Philipp, R., Lindner, W., and Hoffmann, K.-P. (2010). Role of the rostral superior colliculus in gaze anchoring during reach movements. J. Neurophysiol. 103, 3153–3166. https://doi.org/10.1152/jn.00989.2009.

Schroeder, M.R. (1993). Listening with Two Ears. Music Percept. 10, 255–280. https://doi.org/10.2307/40285570.

Schweizer, R., Maier, M., Braun, C., and Birbaumer, N. (2000). Distribution of mislocalizations of tactile stimuli on the fingers of the human hand. Somatosens. Mot. Res. 17, 309–316. https://doi.org/10.1080/08990220020002006.

Sparks, D.L., and Nelson, I.S. (1987). Sensory and motor maps in the mammalian superior colliculus. Trends Neurosci. 10, 312–317. https://doi.org/10.1016/0166-2236(87)90085-3.

Stanford, L.R., and Hartline, P.H. (1984). Spatial and temporal integration in primary trigeminal nucleus of rattlesnake infrared system. J. Neurophysiol. 51, 1077–1090. https://doi.org/10.1152/jn.1984.51.5.1077.

Stein, B.E., and Meredith, M.A. (1993). The merging of the senses. (Cambridge, MA, US: The MIT Press).

Stuphorn, V., Bauswein, E., and Hoffmann, K.P. (2000). Neurons in the primate superior colliculus coding for arm movements in gaze-related coordinates. J. Neurophysiol. 83, 1283–1299. https://doi.org/10.1152/jn.2000.83.3.1283.

Valenstein, E.S. (1965). Independence of approach and escape reactions to electrical stimulation of the brain. J. Comp. Physiol. Psychol. 60, 20–30. https://doi.org/10.1037/h0022299.

Waldbillig, R.J. (1975). Attack, eating, drinking, and gnawing elicited by electrical stimulation of rat mesencephalon and pons. J. Comp. Physiol. Psychol. 89, 200–212. https://doi.org/10.1037/h0076808.

Wall, M.B., Walker, R., and Smith, A.T. (2009). Functional imaging of the human superior colliculus: An optimised approach. Neuroimage 47, 1620–1627. https://doi.org/10.1016/j.neuroimage.2009.05.094.

Wallace, M.T., Wilkinson, L.K., and Stein, B.E. (1996). Representation and integration of multiple sensory inputs in primate superior colliculus. J. Neurophysiol. 76, 1246–1266. https://doi.org/10.1152/jn.1996.76.2.1246.

Weldon, D.A., Calabrese, L.C., and Nicklaus, K.J. (1983). Rotational behavior following cholinergic stimulation of the superior colliculus in rats. Pharmacol. Biochem. Behav. 19, 813–820. https://doi.org/10.1016/0091-3057(83)90086-2.

Werner, W. (1993). Neurons in the Primate Superior Colliculus are Active Before and During Arm Movements to Visual Targets. Eur. J. Neurosci. 5, 335–340. https://doi.org/10.1111/j.1460-9568.1993.tb00501.x.

Werner, W., Dannenberg, S., and Hoffmann, K.P. (1997a). Arm-movementrelated neurons in the primate superior colliculus and underlying reticular formation: Comparison of neuronal activity with EMGs of muscles of the shoulder, arm and trunk during reaching. Exp. Brain Res. 115, 191–205. https://doi.org/10.1007/PL00005690.

Werner, W., Hoffmann, K.P., and Dannenberg, S. (1997b). Anatomical distribution of arm-movement-related neurons in the primate superior colliculus and underlying reticular formation in comparison with visual and saccadic cells. Exp. Brain Res. 115, 206–216. https://doi.org/10.1007/PL00005691.

